# Clinical and primary cell evidence reveals complex CFTR function–phenotype relationships

**DOI:** 10.64898/2026.06.17.732921

**Authors:** Kristen A. Miller, Audrey Pion, Ana Topasna, Alicia J. Ostmann, Jessica D. Meeker, Georgia Kelly, John J. Brewington, Neeraj Sharma, Garry R. Cutting, Karen S. Raraigh

## Abstract

**Rationale:** The CFTR function-phenotype relationship remains incompletely understood, with prior work yielding heterogeneous findings suggesting linear and nonlinear associations.

**Objective:** Define the genotype-function-phenotype relationship using data from the Clinical and Functional TRanslation of *CFTR* (CFTR2) and human nasal epithelial (HNE) studies.

**Methods:** Clinical data (sweat chloride, lung function, pancreatic status) from 84,418 individuals in CFTR2 were linked to CFTR functional measures derived from 289 *CFTR* genotypes. Total genotype function was calculated as the average percent wild-type chloride conductance of both variants in heterologous cell lines. This framework was applied to an HNE cohort including people with CF, CF heterozygotes, and controls. CFTR function was derived from short circuit measurements in HNEs from 153 individuals and correlated with phenotype for 415 individuals. Weighted linear and logarithmic regressions were applied to evaluate the function-phenotype relationship.

**Measurements and Main Results:** Simple linear regression obscured marked heterogeneity across datasets. Piecewise linear regressions revealed marked attenuation of slope magnitude with increasing function across phenotypes. This pattern was well-described by a logarithmic function, such that modeling function on a log scale rendered the relationship approximately linear. HNE data demonstrated similar attenuation, corroborating this pattern.

**Conclusions:** Large-scale natural history data integrated with primary cell findings show that the function-phenotype relationship is not sufficiently described by a single linear effect but is a proportional relationship, in which equivalent changes in CFTR function yield different phenotypic outcomes depending on baseline function. This framework provides precision in predicting clinical benefits from CFTR-directed therapies and identifying meaningful thresholds of CFTR rescue.

**Impact Statement:** This work integrates registry and primary cell data to define the relationship amongst *CFTR* genotype, CFTR protein function, and clinical phenotype. These findings establish reference points for evaluating the degree of phenotypic improvement anticipated from functional restoration from CFTR-targeted treatments. More broadly, this study advances the understanding of CF disease mechanisms by linking molecular function to real-world clinical outcomes across data sources.

**At a Glance Commentary:** *Scientific Knowledge on the Subject:* The relationship between CFTR function and clinical phenotype remains incompletely understood. Prior studies have suggested both linear and nonlinear associations between CFTR activity and disease manifestations. Defining this relationship is increasingly important for interpreting functional data and predicting clinical benefit from CFTR-directed therapies.

*What This Study Adds to the Field:* Using clinical and functional data from more than 84,000 individuals in CFTR2 together with primary human nasal epithelial cell measurements spanning people with cystic fibrosis, carriers, and unaffected controls, we demonstrate that the CFTR function-phenotype relationship is not adequately described by a single linear model. Instead, the relationship is best fitted by piecewise linear regressions of varying slope conforming to a logarithmic pattern, with the greatest phenotypic gains occurring at the lowest levels of baseline CFTR function. These findings provide a quantitative framework for interpreting functional rescue and predicting therapeutic benefit across the CFTR functional spectrum.

## Introduction

Cystic fibrosis (CF) is an autosomal recessive disorder caused by biallelic pathogenic variants in the *CFTR* gene, which encodes the cystic fibrosis transmembrane conductance regulator (CFTR), an epithelial chloride and bicarbonate ion channel(1). Dysfunction of CFTR leads to a multisystem disorder that includes chronic pulmonary disease, exocrine pancreatic insufficiency, gastrointestinal and hepatobiliary manifestations, reproductive tract abnormalities, and elevated sweat chloride concentration (sweat [Cl^-^], typically ≥60 mmol/L)(2).

Clarifying the genotype-function-phenotype relationship is essential for evaluating the level of CFTR function needed for improving or eliminating CF symptoms, particularly in the context of therapeutics development. The substantial diversity in DNA variants (allelic heterogeneity) and corresponding clinical variability amongst individuals with CF provides a unique opportunity to define this relationship across the functional spectrum. While genotype-conferred CFTR function is a major determinant of disease severity, the nature of the relationship between CFTR function and phenotype remains debated. Prior studies support both linear(3, 4) and nonlinear associations(5–8) between CFTR activity and disease traits.

The advent of CFTR modulators – small molecules that act on defective CFTR protein to improve production, intracellular processing, and/or function – has intensified interest in defining these relationships. Highly-effective modulator therapies (HEMT) are now considered standard of care for people with CF (pwCF) with responsive genotypes(9), though a subset has non-modulator responsive genotypes. For these individuals, evaluating the relationship between CFTR function and phenotype will allow assessment of clinical impact and risk/benefit ratio for other interventions, such as gene-based approaches.

Large-scale clinical resources and physiologically relevant model systems offer complementary insights into the nature of the function-phenotype relationship. The Clinical and Functional TRanslation of CFTR (CFTR2) database(10) is a globally-recognized resource that integrates clinical, population-level, and functional data – assays in heterologous cell lines measuring *variant*-specific CFTR function – to interpret the genetic changes identified in pwCF. Human nasal epithelial (HNE) cells obtained via nasal brushing enable the study of *genotype*-specific CFTR function in a native cellular environment. This approach, which determines a person’s baseline or untreated CFTR function in primary nasal cells, may also involve theratyping to enable a personalized approach to CF treatment by testing pharmacological responsiveness of patient-derived respiratory cells(11).

Leveraging expanded CFTR2 data alongside HNE primary cell studies, we sought to delineate the CFTR genotype-function-phenotype relationship. Based on prior work and biological plausibility – including evidence of nonlinear relationships in other monogenic diseases(12–15) – we hypothesize a nonlinear relationship in which small increases in CFTR function at the low end of the functional spectrum correspond to disproportionately large clinical improvements. A precise understanding of this relationship is essential for defining meaningful thresholds for disease amelioration, predicting clinical improvement achievable with therapeutics based on baseline CFTR function, setting realistic expectations regarding anticipated therapeutic benefit, and providing personalized, genotype-specific counseling.

## Methods

### CFTR2 dataset and functional assignment

Genotype and clinical data (sweat [Cl^-^], mean FEV_1_ percent predicted (FEV_1_pp), mean mortality-adjusted Kulich normal residual [KNoRMA] z-scores(16), and pancreatic insufficiency [PI] prevalence) from pwCF were obtained from the CFTR2 database (n=123,005 individuals from 55 countries) as previously described(6). Analysis was restricted to individuals harboring genotypes comprised of exactly two reported *CFTR* variants for which functional characterization was available for both constituent variants and which were reported in at least five people. Excluded genotypes comprised variants classified as non-CF causing by CFTR2 (as of January 2026), deep intronic variants, splice variants, variants known to occur as complex alleles, variants with >75% wild type (WT) function, and variants with no functional measures available. Mean sweat [Cl^-^], mean FEV_1_pp, KNoRMA z-score, and percentage with pancreatic insufficiency (PI) were calculated for genotypes with at least five measurements available.

Variant-specific CFTR function was defined as the average percent WT chloride conductance measured in heterologous cell lines (Fischer rat thyroid [FRT], CF bronchial epithelial [CFBE])(17–19) as previously described. When functional measurements were available from multiple cell types or assays, values were prioritized in the following order: Ussing chamber short-circuit current derived from heterologous CFBE cell lines, Ussing chamber short-circuit current derived from heterologous FRT cell lines, and finally, values resulting from transepithelial chloride conductance (TECC) assays in FRT cells. Variants predicted to result in little-to-no CFTR protein production, referred to as Class I variants(20) (nonsense, canonical splice-site, frameshift, start-loss, and exon deletions/duplications), were grouped as NULL and assigned 0% CFTR activity, though variants perceived as NULL but which may allow CFTR protein production (exceptional Class I variants) were excluded(21). For each genotype, total CFTR function was calculated as the mean percent WT conductance across both alleles.

### HNE dataset and functional assignment

An independent cohort included individuals with a clinical diagnosis of CF for whom human nasal epithelial (HNE) functional measurements and clinical data (sweat [Cl⁻] and/or FEV_1_pp) were available. Clinical values were paired with corresponding genotype-level CFTR functional values obtained in patient-derived HNEs using previously described approaches(5, 22).

To further evaluate the full functional continuum, additional data was incorporated from unaffected adult CF heterozygotes (either F508del/WT or NULL/WT) and asymptomatic, non-heterozygous controls (WT/WT). An accompanying manuscript (Pion, et al.) includes functional testing of HNE cells from CF heterozygotes (n=52) and controls (n=24). These values were linked with sweat [Cl^-^] measurements from previously published literature (n=47 CF heterozygotes and n=291 controls) summarized in **Supplemental Tables 1 and 2**. Functional values for CF heterozygotes were expressed as percent WT function relative to asymptomatic controls. Data sources for the full HNE cohort (pwCF, CF heterozygotes, and controls) are summarized in **Table 1**.

**Table 1.**
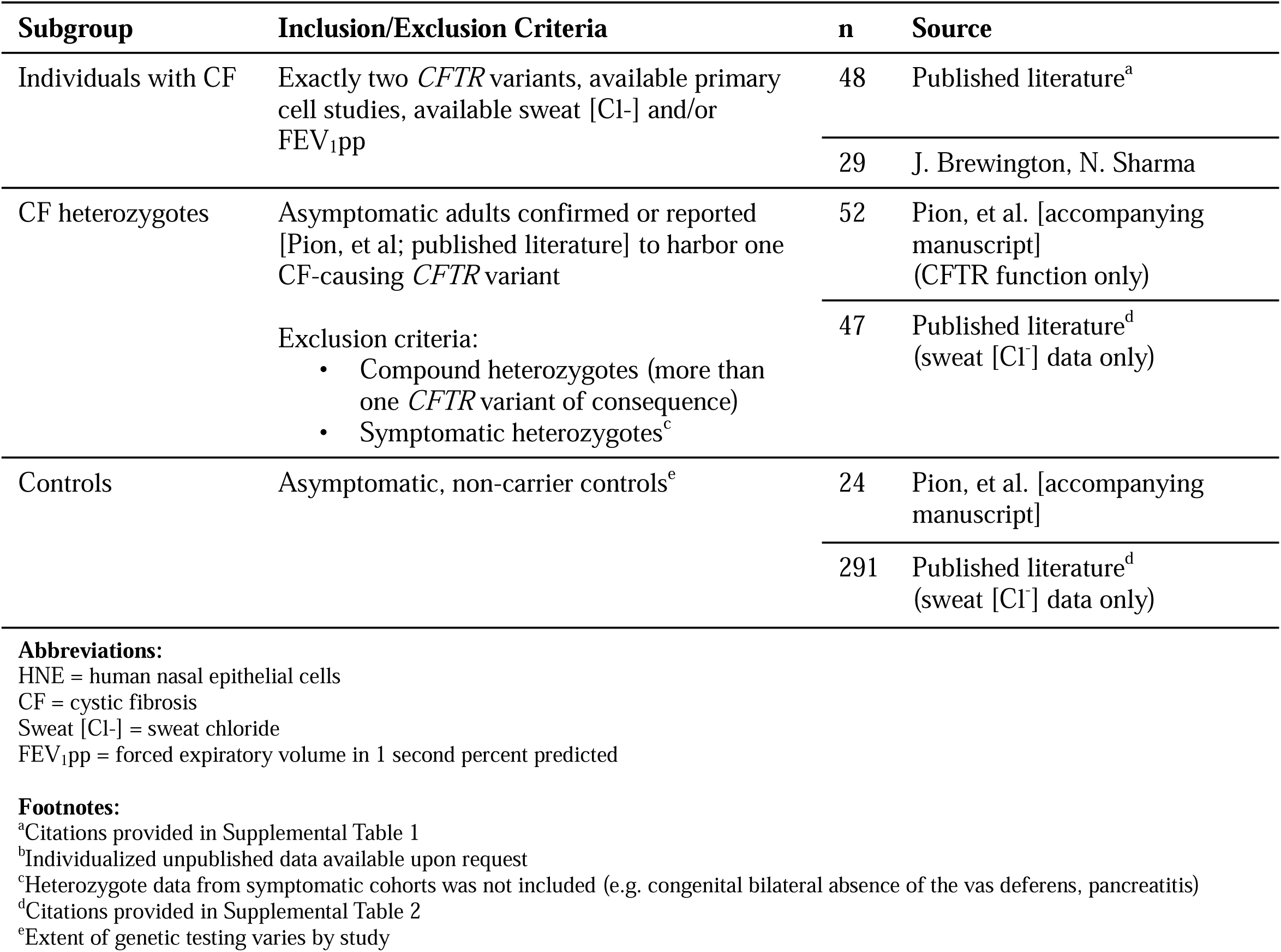
Sources of HNE function and associated clinical data.

### Statistical analysis

Analyses were performed separately in each cohort (CFTR2 and HNE) using linear regression models weighted by the number of individuals contributing to phenotypic measures (CFTR2) or functional measures (HNE) for each genotype-level estimate where applicable. Initial models evaluated a single linear relationship between CFTR function and clinical phenotype across the full functional range. To explore potential deviation from linearity, piecewise linear regression models allowing for multiple slopes across specified functional ranges were constructed. Secondary analyses on sweat [Cl^-^] data for both CFTR2 and HNEs evaluated logarithmic models. Statistical analyses were performed using Stata version 15.0.

## Results

### Patient makeup and functional distribution of 289 CFTR genotypes from CFTR2

A total of 84,418 individuals from the CFTR2 dataset harbored one of 289 genotypes present in at least five individuals and for which function could be assigned (**Figure 1A**). Genotype function ranged from 0% to 37.25% in pwCF; 94% of genotypes fell into the 0-10% range (**Figure 1B**).

**Figure 1.**
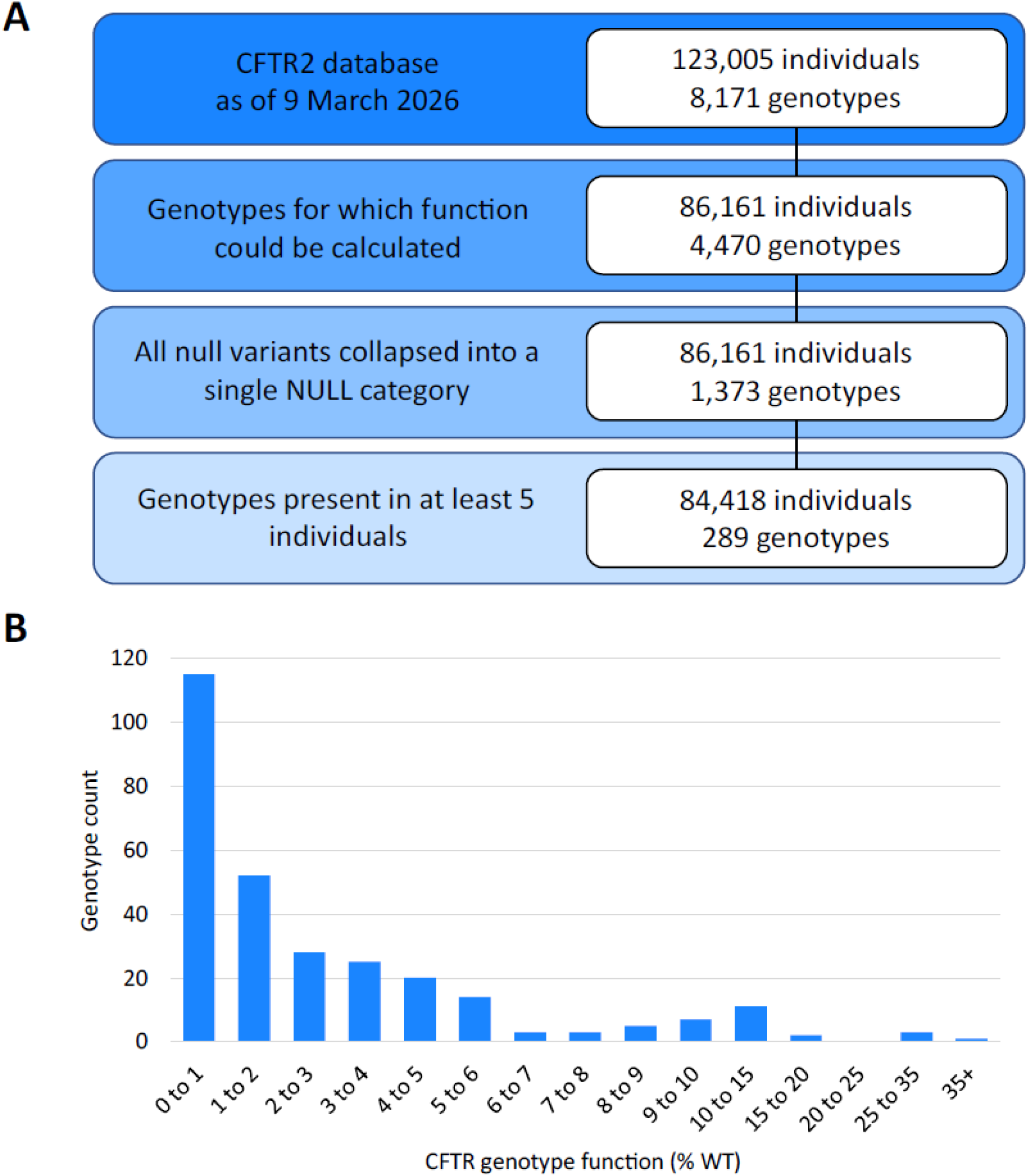
Inclusion flow diagram and functional categorization of genotypes from individuals in CFTR2 analyzed for function – phenotype relationships. **A** The dataset from the Clinical and Functional Translation of CFTR (CFTR2) project comprised 8,171 genotypes across 123,005 individuals, contributed by participating registries and cystic fibrosis centers. Genotypes containing a single null variant were grouped so that any null allele could occur in *trans* with a non-null variant of interest. Genotypes with two null variants were consolidated into a single null/null category. Only genotype groups present in at least five individuals were retained, yielding 289 genotypes harbored by 84,418 for analysis. B The distribution of genotype function across the CFTR2 dataset revealed that the vast majority of individuals included in analysis had function <10% of WT, consistent with CF. Genotype function, expressed as percent of wild-type activity (%WT), was calculated as the mean functional value of the two variants comprising each genotype. **Alt Text:** Flow diagram showing selection of 289 CFTR genotypes from the CFTR2 database and histogram demonstrating that most analyzed genotypes have less than 10% wild-type CFTR function.

### Sweat [Cl^-^] evaluation using CFTR2 data

Sweat [Cl^-^] was the first trait for evaluation of the CFTR genotype-function-phenotype because it is highly correlated with CFTR function and is not significantly impacted by modifier genes or environmental factors(23). While use of heterologous cell-based systems has proven useful in defining CFTR function and aiding in the determination of disease liability, discrimination at very low levels of residual function is poor due to technical limitations(24). Indeed, the magnitude of change amongst genotypes with 0 to 1% of WT CFTR function, although statistically significant (slope = -2.1, root mean square error [RMSE] 1.4 p<0.001), was not considered clinically meaningful (**Supplemental Figure 1**). Therefore, all genotypes with function <1% were grouped as 1% functional activity for subsequent analyses.

Upon grouping all genotypes with less than 1% CFTR function and plotting the remaining data, function derived from unique genotypes ranged from 1 to 37.5% and mean genotype-derived sweat [Cl^-^] ranged from 39 to 119 mmol/L (**Figure 2, panel A**). While a global linear model demonstrated a significant (p<0.001) inverse association between CFTR function and sweat [Cl-] in CFTR2 data (slope = -4.5, RMSE 5.3), this approach obscured substantial heterogeneity (**Figure 2, panel B**). Therefore, biologically relevant thresholds were chosen to evaluate whether the relationship differed across the functional spectrum. Weighted piecewise linear regression models were applied to genotypes with function <10% of WT function (associated with development of clinical CF disease) and ≥10% (associated with reduced or non-penetrance for CF) and demonstrated markedly different slopes (<10%: slope = -7.9, RMSE 4.2; ≥10%: slope = -0.7, RMSE 8.1) and an improved RMSE for <10% function when compared to the single linear model (**Figure 2C**). This indicates substantially greater phenotypic change per unit of CFTR function at lower functional levels. A second biologically-relevant threshold of 5% was also applied, reflecting the boundary below which pancreatic insufficiency is typical and above which pancreatic sufficiency or attenuated disease may occur(25). Weighted piecewise linear regression using these two thresholds revealed marked attenuation of slope magnitude with increasing CFTR function, with the steepest relationships observed at low baseline function (<5%: slope = -10.1, RMSE 3.7) and diminished phenotypic change at higher levels of baseline function (5-10%: slope = -2.5, RMSE 8.9; ≥10%: slope = -0.7, RMSE 8.1; **Figure 2, panel D**). Collectively, this pattern suggested a nonlinear relationship, leading to further regression modeling. Both the fitted logarithmic model (**Figure 2, panel E**) and semi-log visualization (**Figure 2, panel F**) effectively represented the interaction between the variables, as indicated by the low RMSE and improved visual fit. Consequently, the three-segment weighted piecewise model (specifically the <5% segment) and logarithmic model were selected for application to additional CF disease traits.

**Figure 2.**
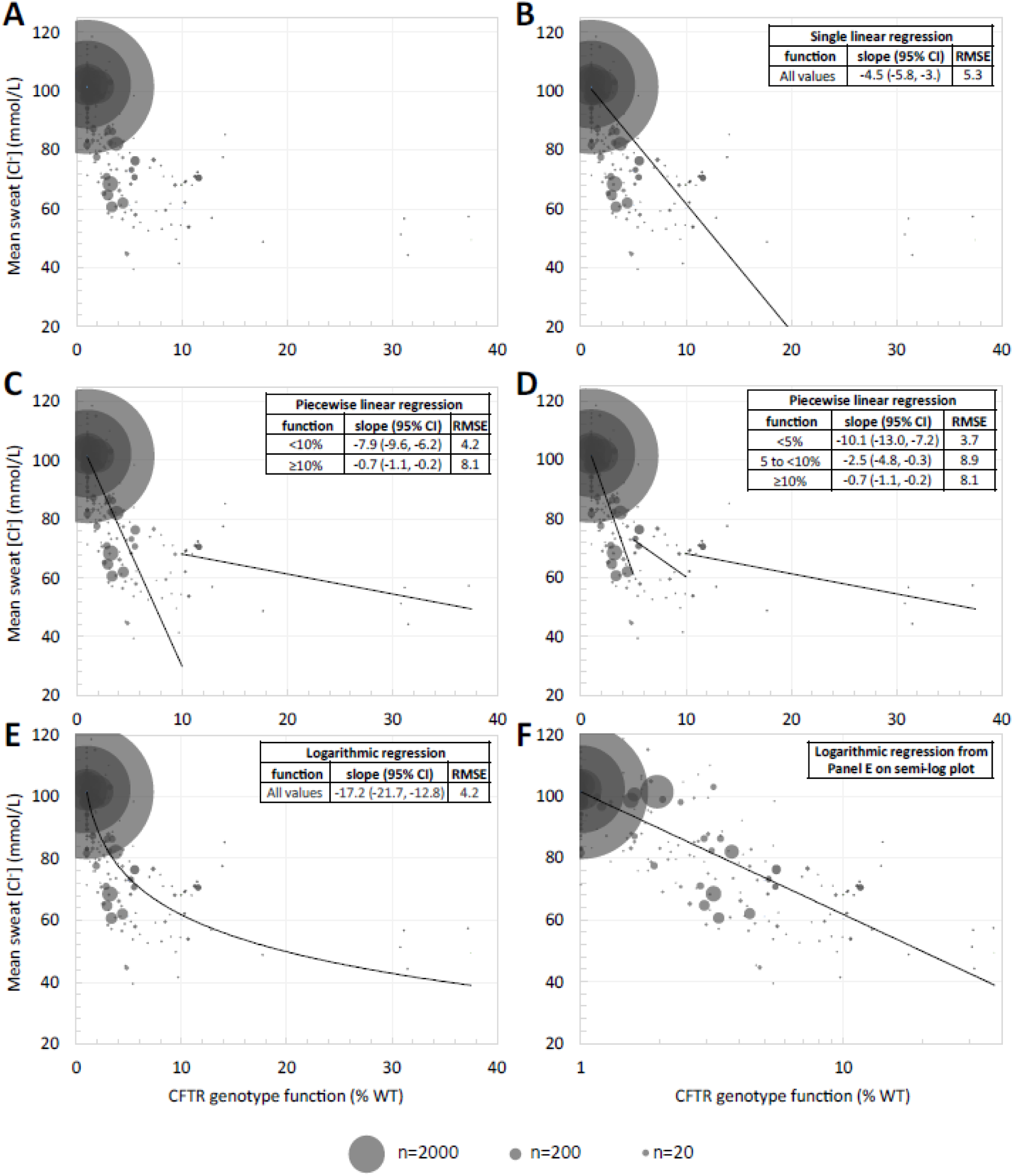
Analysis of the function-phenotype relationship in the CFTR2 dataset using single linear, piecewise linear, and logarithmic regressions to evaluate sweat chloride concentration. The relationship between *CFTR* genotype function (expressed as percentage of wild-type [% WT] chloride conductance) and mean sweat chloride concentration [Cl^-^] is shown using either linear (y=*m*x+*b*) or logarithmic (y=*m*ln(x)+*b*) regressions. Each bubble represents a genotype-level mean, with bubble size reflecting the number of sweat [Cl^-^] measurements contributing. The largest bubbles correspond to common genotypes: F508del/F508del and F508del/NULL. Regressions are weighted by sweat [Cl^-^] n. Slopes and root mean square error (RMSE) for each model across specified functional ranges are reported in the inlay tables. **A** Data shown with no regression analysis. **B** Data shown with a single linear model across the full functional range of values. **C** Data shown with piecewise linear regression, segmented at the biologically-relevant threshold of 10% CFTR function, below which a CF phenotype is expected and above which the penetrance for CF is reduced. **D** Data shown with piecewise linear regression, further segmented at the biologically-relevant threshold of 5% CFTR function, below which pancreatic insufficient CF (PI-CF) is expected, and at 10% as described in panel C. Individuals with CF who have CFTR function 5-10% are likely to be pancreatic sufficient. **E** Data shown with a logarithmic model as an example of a single function producing a curve to describe the data. **F** Data shown from panel E on a semi-log plot; when the x-axis is a logarithmic scale, the data becomes approximately linear. **Alt Text:** Scatterplots showing an inverse relationship between CFTR function and sweat chloride in CFTR2. Piecewise and logarithmic models fit the data better than a single linear regression.

### Pancreatic insufficiency prevalence, FEV_1_pp, and KNoRMA z-score evaluation using CFTR2 data

The relationship between CFTR function and phenotype was significant for all clinical traits analyzed (pancreatic insufficiency [PI] prevalence, FEV_1_pp, and KNoRMA z-score). A single linear regression model is shown for all traits (**Figure 3, panels A, D, and G**, **dashed line**) and, while significant (p<0.001 for all traits), again obscured heterogeneity across the functional spectrum. Weighted piecewise linear regression models fit at <5%, 5-10%, and ≥10% revealed a similar pattern to sweat [Cl^-^]: continuously attenuated slope magnitude with increasing function, with PI prevalence having the most significant changes across thresholds (<5%: slope = -22.0, RMSE 10.4; 5-10%: slope = -0.4, RMSE 17.0; ≥10%: slope = -0.1, RMSE 17.2). Fitting to a logarithmic model improved RMSE over the global linear model for all traits (**Figure 3, panels A-I**).

**Figure 3.**
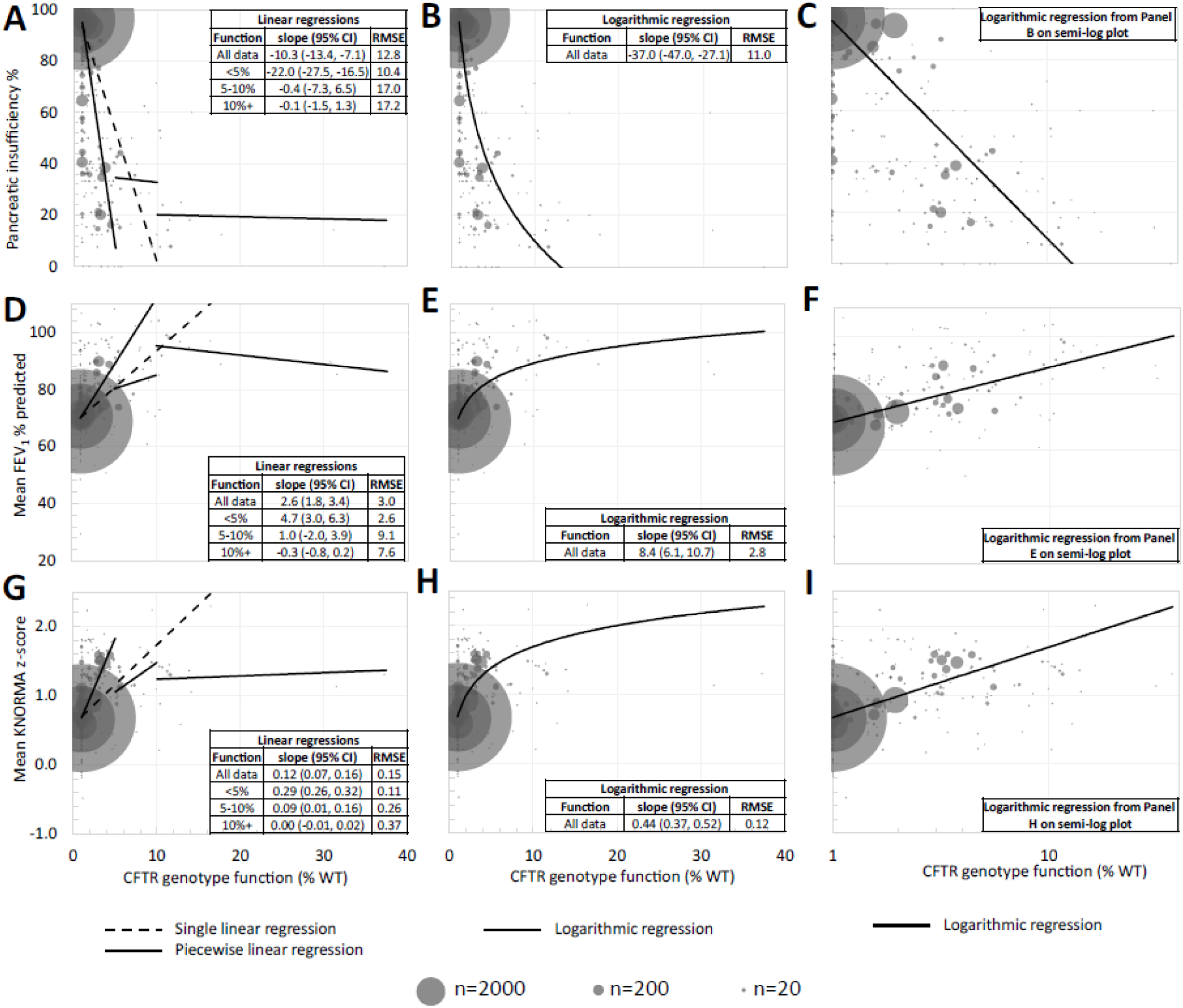
Analysis of the function-phenotype relationship in the CFTR2 dataset using linear and logarithmic regression to evaluate pancreatic insufficiency prevalence, FEV1 % predicted, and KNORMA z-score. The relationship between *CFTR* genotype function (expressed as percentage of wild-type [% WT] chloride conductance) and pancreatic insufficiency prevalence **(top row, A-C)**, mean FEV1 % predicted **(middle row, D-F)**, and mean KNoRMA z-score **(bottom row, G-I)** is shown using linear (y=*m*x+*b*) and logarithmic (y=*m*ln(x)+*b*) regressions. Each bubble represents a genotype-level mean, with bubble size reflecting the number of measurements contributing. The largest bubbles correspond to common genotypes: F508del/F508del and F508del/NULL. Regressions are weighted by the number of measures contributing to a datapoint. Slopes and root mean square error (RMSE) for each model across specified functional ranges are reported in the inlay tables. **First Column (A, D, G)** Data shown with piecewise linear regression, segmented at the biologically-relevant thresholds of 5% and 10% CFTR function (corresponding to approximate thresholds for pancreatic sufficiency and reduced penetrance, respectively). **Middle Column (B, E, H)** Data shown with a logarithmic model as an example of a single function producing a curve to describe the data. **Last Column (C, F, I)** Data shown from panel B on a semi-log plot; when the x-axis is a logarithmic scale, the data becomes approximately linear. **Alt Text:** Graphs showing relationships between CFTR function and pancreatic insufficiency, FEV_1_pp, and KNoRMA score. Phenotypic changes are greatest at low CFTR function and attenuate at higher function.

### Phenotype and functional distribution of HNE cohort

To examine the relationship between function and phenotype (sweat [Cl^-^] and FEV_1_pp) in primary cells, an independent HNE cohort representing 45 *CFTR* genotypes was assembled using data from individuals with CF (n=77 individuals harboring 42 genotypes), heterozygous CF heterozygotes (n=52 individuals with one copy of F508del or one NULL variant who underwent functional testing), and asymptomatic controls (n=24 individuals who underwent functional testing) compiled from published datasets and investigator-contributed cohorts (**Table 1**). Inclusion of CF heterozygotes and controls substantially increased representation at higher functional levels, which were sparsely represented in the CFTR2 patient base.

### Sweat [Cl^-^] evaluation using HNE data

Functional values derived from HNE studies and sweat [Cl^-^] measures were plotted and demonstrated a range of values (**Figure 4, panel A**). Due to the significantly smaller dataset compared to CFTR2, individual measurements were plotted with each dot representing a single person, with the exception of CF heterozygotes and controls, which represent composite functional and clinical data. Regressions were weighted by the number of individuals who contributed measurements to the mean functional values (relevant for CF heterozygotes and controls only). A weighted global linear model demonstrated significant association between CFTR function and sweat [Cl□] (slope = -0.9, RMSE 17.1, p<0.001) but oversimplified the dispersion of the data as observed with the CFTR2 dataset (**Figure 4, panel B**). Weighted piecewise linear regression at biologically-relevant thresholds again showed attenuation of slope magnitude at function >10% (**Figure 4, panels C and D**). Fitting the data to a single weighted logarithmic model demonstrated improvement in RMSE over the weighted single linear model (12.7 vs. 17.1; **Figure 4, panels E and F**).

**Figure 4.**
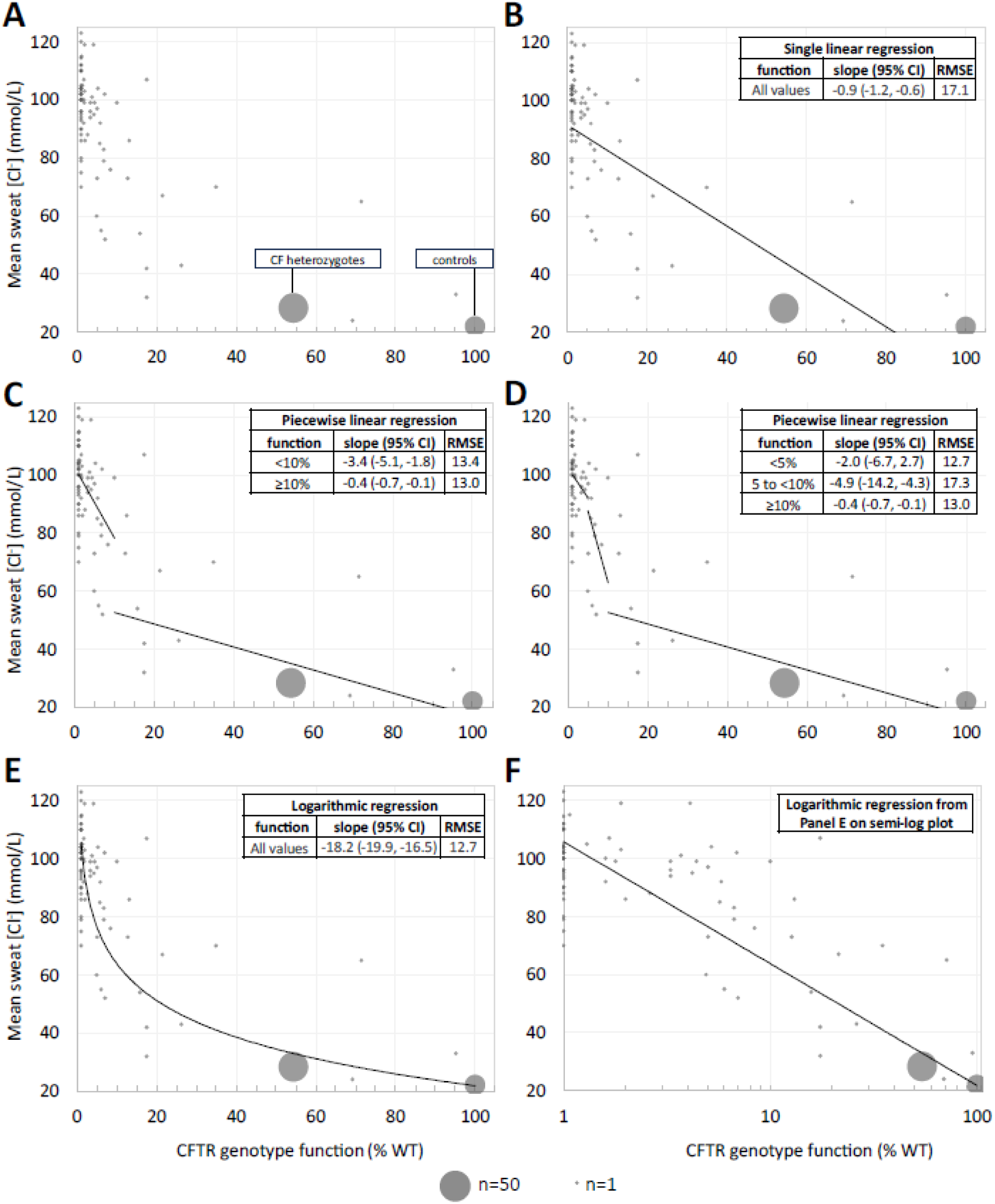
Analysis of the function-phenotype relationship in primary HNE cells using linear and logarithmic regression. The relationship between *CFTR* genotype function (expressed as percentage of wild-type [% WT] chloride conductance) and sweat chloride concentration [Cl^-^] is shown using either linear (y=*m*x+*b*) or logarithmic (y=*m*ln(x)+*b*) regressions. Each point represents a single person whose CFTR function and sweat [Cl^-^] are plotted, with the exception of heterozygous CF heterozygotes (largest bubble, n=52 individuals whose CFTR function was measured and aggregated to a mean value of 54.2% of WT-CFTR function) and healthy non-carrier controls (second-largest bubble, n=24 individuals whose CFTR function was measured and the mean value set to 100% of WT-CFTR function). Regressions are weighted by the number of functional measures per bubble. Slopes and root mean square error (RMSE) for each model across specified functional ranges are reported in the inlay tables. **A** Data shown with no regression analysis. **B** Data shown with a single linear model across the full functional range of values. **C** Data shown with piecewise linear regression, segmented at the biologically-relevant threshold of 10% CFTR function, below which a CF phenotype is expected and above which the penetrance for CF is reduced. **D** Data shown with piecewise linear regression, further segmented at the biologically-relevant threshold of 5% CFTR function, below which pancreatic insufficient CF (PI-CF) is expected, and at 10% as described in panel C. Individuals with CF who have CFTR function 5-10% are likely to be pancreatic sufficient. **E** Data shown with a logarithmic model as an example of a single function producing a curve to describe the data. **F** Data shown from panel E on a semi-log plot; when the x-axis is a logarithmic scale, the data becomes approximately linear. **Alt Text:** HNE data showing an inverse relationship between CFTR function and sweat chloride across individuals with cystic fibrosis, CF heterozygotes, and controls. Logarithmic modeling best describes the relationship.

### FEV_1_pp evaluation using HNE data

The same framework of plotting HNE-derived functional measures and CF phenotype data was applied for FEV_1_pp. Because no FEV_1_pp measures were available for CF heterozygotes and controls, each data point represents n=1 pwCF and regressions are unweighted. A single linear regression (dashed line) as well as piecewise regressions (solid lines) are shown in **Figure 5, panel A**. No regression demonstrated a significant relationship at any functional range (p>0.05 for all analyses) despite visually appearing to follow a similar trend to other data in which a steeper slope is observed at <5% of WT CFTR function compared to >10%. Fitting the data to a logarithmic model (**Figure 5, panels B and C**) did not significantly improve RMSE and did not visually differ significantly from the global linear model, with significant dispersion in data points along the functional continuum.

**Figure 5.**
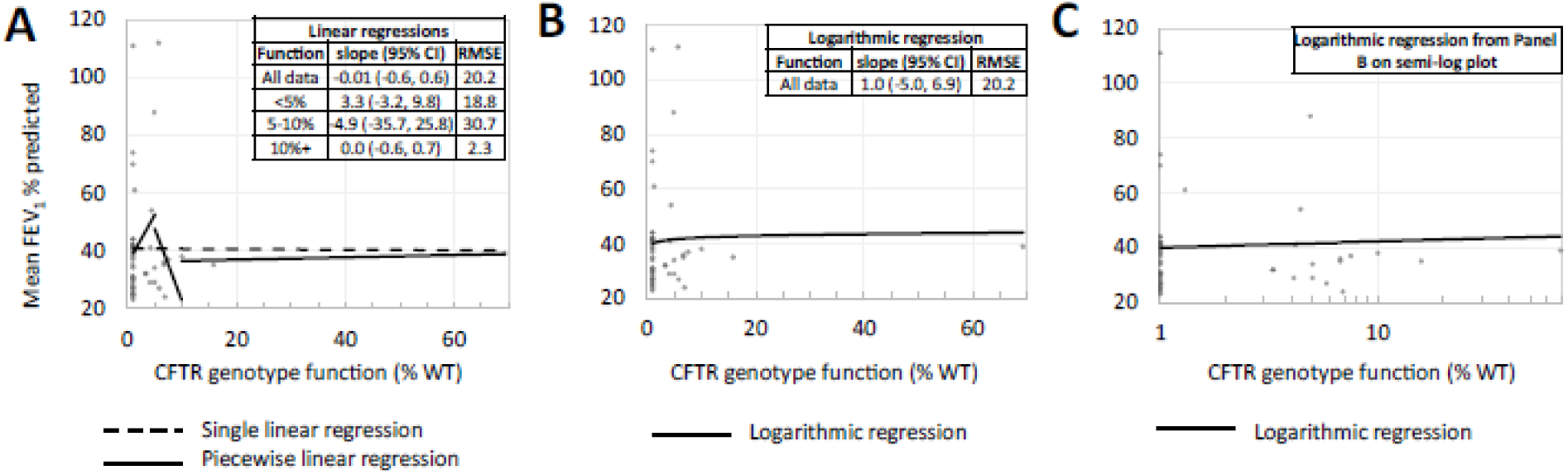
Analysis of the function-phenotype relationship in primary HNE cells linear and logarithmic regression to evaluate FEV_1_ % predicted. The relationship between *CFTR* genotype function (expressed as percentage of wild-type [% WT] chloride conductance) and FEV_1_ % predicted is shown using linear (y=*m*x+*b*) and logarithmic (y=*m*ln(x)+*b*) regressions. Each bubble represents n=1 person who underwent CFTR functional testing and had a recorded cross-sectional FEV_1_ % predicted value. Slopes and root mean square error (RMSE) for each model across specified functional ranges are reported in the table. No significant association between CFTR function and FEV % predicted was identified for the global linear regression, any piecewise regression segment, or the logarithmic model (all p > 0.05). **A** Data shown with a single linear and piecewise linear regression, segmented at the biologically-relevant thresholds of 5% and 10% CFTR function (corresponding to approximate thresholds for pancreatic sufficiency and reduced penetrance, respectively). **B** Data shown with a logarithmic model as an example of a single function producing a curve to describe the data. **C** Data shown from panel B on a semi-log plot; when the x-axis is a logarithmic scale, the data becomes approximately linear. **Alt Text:** Graphs evaluating CFTR function and FEV1 percent predicted in human nasal epithelial cell studies. Although trends resemble other phenotypes, no statistically significant association was observed.

### Comparison of the function-phenotype relationship using CFTR2 and HNE data

Given the similar shapes of the piecewise regressions and logarithmic models across both CFTR2 and HNE datasets in evaluation of the relationship between CFTR function and sweat [Cl^-^], we assessed whether the models derived in two independent datasets were comparable to each other. The weighted single linear regressions for each dataset do not align, with slopes of - 4.5 (CFTR2 data) and -0.9 (HNE data; **Figure 6, panel A**). However, the weighted logarithmic models in each dataset have nearly-complete overlap across the functional spectrum and essentially equal slopes of -17.2 and -18.2 (CFTR2 and HNE data, respectively; **Figure 6, panel B**). Displaying the data on a semi-log plot demonstrated better visualization of the data and approximates linearity, consistent with a logarithmic function being a reasonable model to use (**Figure 6, panel C**).

**Figure 6.**
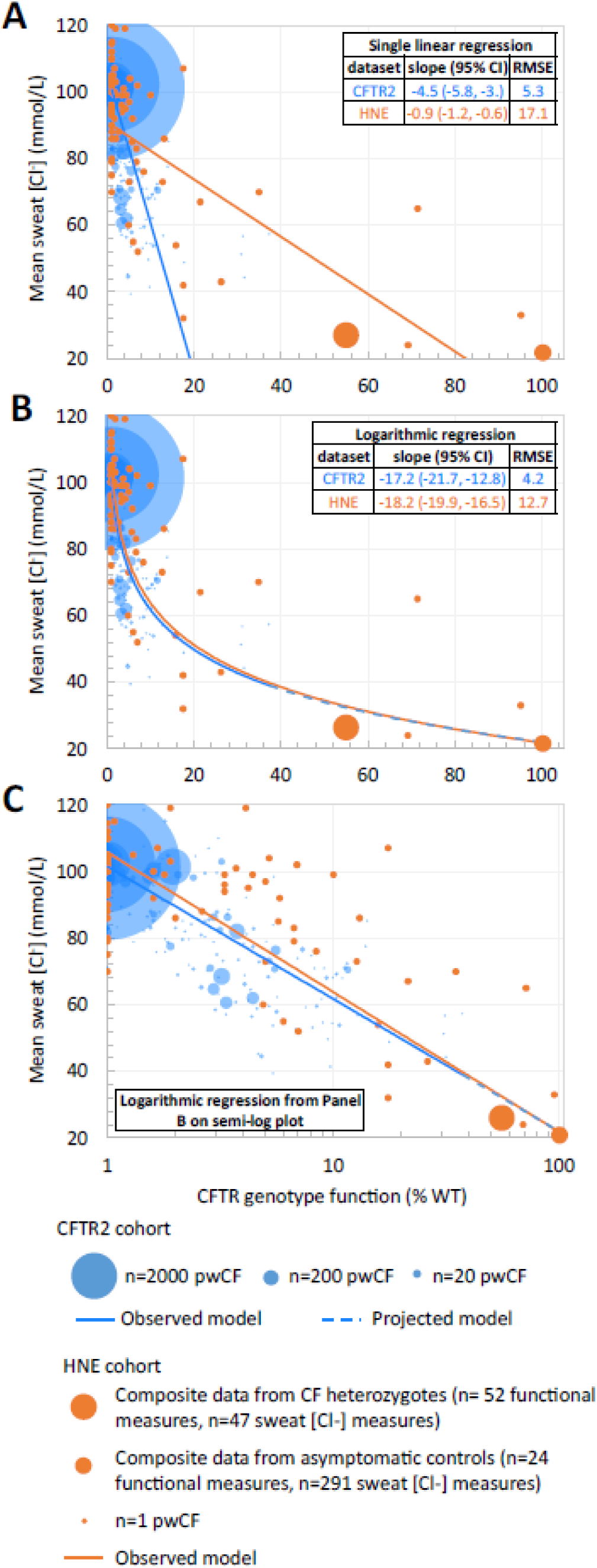
The function – phenotype relationship as determined using CFTR2 and primary HNE data can be modeled as a single logarithmic curve. **A** Data derived from CFTR2 is shown in blue. Each bubble represents a genotype-level mean, with bubble size reflecting the number of measurements contributing to that estimate. The largest bubbles correspond to common genotypes: F508del/F508del and F508del/NULL. Data derived from HNE studies is shown in orange. Each circular data point represents 1 individual with CF. Larger orange circles indicate either unaffected heterozygote CF heterozygotes or unaffected, non-carrier controls from whom CFTR functional measures were derived using HNE cells while sweat [Cl^-^] measures were derived from the published literature. **B** Logarithmic regressions (weighted by sweat [Cl^-^] n for CFTR2 data) were fit across the entire functional spectrum, with slope and intercepts reported in the inlay table. **C** The same data from panel B are shown on a semi-log plot, with logarithmic regressions now appearing linear when the x-axis scale is logged. **Alt Text:** Comparison of CFTR2 and HNE datasets showing similar logarithmic relationships between CFTR function and sweat chloride despite differing linear regression slopes.

## Discussion

Measures of CFTR activity via nasal potential difference or in response to CFTR modulator treatment have been interpreted as demonstrating an approximately linear relationship(3, 4, 26). However, several prior studies have visually suggested nonlinear relationships between CFTR function and disease manifestations, particularly for sweat [Cl-]. In addition to McCague et al.(6), Brewington and colleagues demonstrated a nonlinear inverse relationship between CFTR-mediated short-circuit current and sweat [Cl-] in primary epithelial cell models(5), and Char et al. similarly observed nonlinearity between CFTR function and sweat [Cl-] measurements(7). Work examining acquired CFTR dysfunction in cigarette smoke exposure has likewise suggested attenuation of phenotypic effects at higher levels of CFTR activity across the spectrum of asymptomatic controls, smokers, and people with CF(27).

Our study leverages natural history registry data and functional characterization from heterologous cell lines, together with a primary cell HNE cohort data, to refine the relationship between CFTR genotype function and clinical phenotypes. A prior analysis of CFTR2 data demonstrated a logarithmic relationship between CFTR function and sweat [Cl-], pancreatic status, and lung function(6), and a recent real-world study similarly reported that sweat [Cl^-^] response to elexacaftor/tezacaftor/ivacaftor was only slightly greater in individuals with two vs. one responsive variants(28), consistent with near-saturation of phenotypic effect at higher functional levels achieved from treatment. While these models support nonlinearity, they do not fully characterize the way in which the slope may vary across the functional or phenotypic spectrum.

Using expanded CFTR2 data with broader genotype representation supplemented by primary cell HNE data extending into carrier and WT functional ranges, our findings demonstrate important phenotype- and function-specific heterogeneity not captured by a single model. Models restricted to narrower functional ranges revealed marked differences in slope magnitude using piecewise linear regression. Visualizing CFTR function on a logarithmic axis produced a more consistent linear relationship across the observed range – findings also corroborated in the independent HNE cohort. As expected, phenotypes such as FEV_1_pp demonstrated broader variability, likely reflecting the influence of non-CFTR factors including genetic modifiers, environmental exposures, and disease progression(16, 29). Nevertheless, the persistence of consistent directional trends across phenotypes – including those associated with morbidity and mortality – supports an underlying nonlinear relationship between CFTR function and clinical expression. The steep decline in PI prevalence approaching 5% function, followed by a marked plateau above this threshold, reinforces this level as an important inflection point in disease expression and a clinically relevant therapeutic target. Together, these results indicate that the relationship between CFTR function and phenotype is not adequately described by a single linear model but instead reflects differential phenotypic impact of incremental changes in CFTR activity across the functional spectrum, with possible contributions from tissue-specific regulation, nonlinear ion transport dynamics, and genetic or environmental modifiers.

An important strength of the present study is the complementary primary cell data, including incorporation of CF heterozygotes and unaffected controls into the functional spectrum analyzed. While a recent report found ∼80% function in human bronchial epithelial cells of CF heterozygotes(4), the data from the accompanying manuscript by Pion et al. included in our study consistently approximated 50% function in CF heterozygotes, which aligns with expectations for a classic additive model for traditional recessive inheritance. This distinction has important implications for interpreting the function-phenotype relationship across the entire continuum, as carrier and control values provide an important biologic anchor beyond the severely reduced functional ranges represented in most CF cohorts. Differences between studies could reflect a variety of factors, including differences in assay systems, normalization approaches, study populations, or tissue-specific biology. If heterozygotes retain CFTR activity closer to unaffected controls, the functional reserve separating unaffected individuals from CF disease states would be substantially widened. In contrast, our findings support the presence of an intermediate functional state in CF heterozygotes, with mean CFTR activity approximately half that observed in unaffected controls, reinforcing the concept that phenotypic consequences of incremental changes in CFTR activity are not uniform across the functional spectrum.

Understanding the relationship between CFTR function and clinical phenotype is central to the development and evaluation of CFTR-directed therapies. This framework may be particularly relevant for evaluating emerging therapies, including modulators in very low-function genotypes and non-modulator approaches such as gene-based therapies, where anticipated gains in CFTR activity must be translated into expected clinical benefit. Projected levels of functional rescue for methods such as gene replacement, gene editing, RNA-based therapies, or engineered cellular therapies may be modest, or may vary considerably across platforms and tissues. Preclinical gene editing studies have reported CFTR functional restoration ranging from 5 to >50% of wild-type levels depending on the approach and cellular context(30–32), and in *vivo* delivery constraints are likely to place real-world restoration toward the lower end of this range. Our findings suggest that even small increases in function have the potential to produce a significant reduction in disease burden, particularly in individuals with very low baseline function, where the logarithmic relationship forecasts the steepest phenotypic gains per unit of functional rescue. As non-modulator therapies advance, the nonlinear function–phenotype framework described here may help guide expectations for clinical efficacy and have practical implications for trial design and endpoint selection. A meaningful proportion of pwCF remains ineligible for or intolerant of current CFTR modulators(33), and for these individuals, even partial functional restoration through gene-based approaches could confer substantial clinical benefit if the logarithmic relationship holds in *vivo*.

Several factors should be considered when interpreting the findings from this study. CFTR functional estimates applied to the CFTR2 data were assigned at the genotype level and derived primarily from heterologous expression systems in which individual CFTR alleles are studied in isolation, an approach that may not fully capture the combined effects of both alleles or tissue-specific CFTR activity in *vivo*. Registry data is subject to measurement variability and differences in ascertainment across clinical centers. As expected for a disease database, the majority of genotypes represented in the dataset were associated with very low levels of CFTR function, resulting in comparatively sparser data at higher functional levels. In addition, inter-individual variability related to factors such as age, environmental exposures, and genetic modifiers may influence clinical phenotype independently of *CFTR* genotype and are not accounted for in the present analysis. In the HNE cohort, data were derived from multiple sources, including published literature and assays performed across different laboratories, which may introduce additional heterogeneity. For most of the cohort, the derived CFTR function values and sweat [Cl□] data represented paired measurements from the same individuals. For CF heterozygotes and controls, however, CFTR functional measures were obtained in a single laboratory, while sweat [Cl□] data were drawn from multiple published reports. Finally, the primary cell HNE cohort included a smaller number of genotypes and participants compared to CFTR2, limiting precision of estimates derived from these data.

Future work may extend these analyses to additional CF phenotypes such as infection or CF-related diabetes to determine whether similar nonlinear relationships are observed across other disease traits. Further refinement of models, including comparison of alternative nonlinear model forms, may help identify the most accurate representation of CFTR function–phenotype relationships. Clinical trial datasets may also allow direct comparison of observed changes in sweat [Cl^-^] following treatment with those expected based on estimated increases in CFTR function associated with HEMTs.

## Conclusion

The relationship between CFTR function and clinical phenotype is not adequately described by a single linear model but instead reflects differential phenotypic impact across the functional spectrum that is described by piecewise linear regression or a logarithmic relationship. A quantitative framework that accounts for this nonlinearity improves interpretation of functional evidence, enhances precision medicine in CF through genotype-informed counseling and accurate prediction of therapeutic response, and guides evaluation of emerging treatments aimed at partial restoration of CFTR activity.

## Author Contributions

K.A.M., K.S.R., and G.R.C. were responsible for conception/design. All authors contributed to data acquisition, data analysis, data interpretation, drafting/reviewing critically for important intellectual content, and approval for publication.

## Online Data Supplement

This article has an online data supplement.

## Data availability statement

Data from the Clinical and Functional Translation of CFTR (CFTR2) cohort is available in the online supplement. Functional and clinical data from human nasal epithelial cell studies was compiled from published literature cited in the supplement, and from investigator-contributed data, which is available upon reasonable request and subject to applicable data sharing and privacy considerations.

## Ethical approval statement

Data obtained from the Clinical and Functional Translation of CFTR (CFTR2) project were collected under JHH IRB NA_00018599. Human nasal epithelial studies were conducted under JHH IRB 00116966 and CCHMC IRB# 2015-2757.

## Acknowledgements

The authors acknowledge the Clinical and Functional Translation of CFTR (CFTR2) project for providing access to genotype, clinical, and functional data used in this study. We are grateful to the individuals with cystic fibrosis, their families, participating clinical centers, national registries, and the CFTR2 investigators whose contributions have made this resource possible.

## References

1. Cutting GR. Cystic fibrosis genetics: from molecular understanding to clinical application. Nat Rev Genet 2015; 16: 45–56.

2. Grasemann H, Ratjen F. Cystic Fibrosis. N Engl J Med 2023; 389: 1693–1707.

3. Accurso FJ, Van Goor F, Zha J, Stone AJ, Dong Q, Ordonez CL, Rowe SM, Clancy JP, Konstan MW, Hoch HE, Heltshe SL, Ramsey BW, Campbell PW, Ashlock MA. Sweat chloride as a biomarker of CFTR activity: proof of concept and ivacaftor clinical trial data. J Cyst Fibros 2014; 13: 139–147.

4. Zemanick ET, Ramsey B, Sands D, McKone EF, Fajac I, Taylor-Cousar JL, Mall MA, Konstan MW, Nair N, Zhu J, Arteaga-Solis E, Van Goor F, McGarry L, Prieto-Centurion V, Sosnay PR, Bozic C, Waltz D, Mayer-Hamblett N. Sweat chloride reflects CFTR function and correlates with clinical outcomes following CFTR modulator treatment. J Cyst Fibros 2025; 24: 246–254.

5. Brewington JJ, Filbrandt ET, LaRosa FJ, Moncivaiz JD, Ostmann AJ, Strecker LM, Clancy JP. Brushed nasal epithelial cells are a surrogate for bronchial epithelial CFTR studies. JCI Insight 2018; 3.

6. McCague AF, Raraigh KS, Pellicore MJ, Davis-Marcisak EF, Evans TA, Han ST, Lu Z, Joynt AT, Sharma N, Castellani C, Collaco JM, Corey M, Lewis MH, Penland CM, Rommens JM, Stephenson AL, Sosnay PR, Cutting GR. Correlating Cystic Fibrosis Transmembrane Conductance Regulator Function with Clinical Features to Inform Precision Treatment of Cystic Fibrosis. Am J Respir Crit Care Med 2019; 199: 1116–1126.

7. Char JE, Wolfe MH, Cho HJ, Park IH, Jeong JH, Frisbee E, Dunn C, Davies Z, Milla C, Moss RB, Thomas EA, Wine JJ. A little CFTR goes a long way: CFTR-dependent sweat secretion from G551D and R117H-5T cystic fibrosis subjects taking ivacaftor. PLoS One 2014; 9: e88564.

8. Rowe SM, Accurso F, Clancy JP. Detection of cystic fibrosis transmembrane conductance regulator activity in early-phase clinical trials. Proc Am Thorac Soc 2007; 4: 387–398.

9. Taylor-Cousar JL, Robinson PD, Shteinberg M, Downey DG. CFTR modulator therapy: transforming the landscape of clinical care in cystic fibrosis. Lancet 2023; 402: 1171–1184.

10. Sosnay PR, Siklosi KR, Van Goor F, Kaniecki K, Yu H, Sharma N, Ramalho AS, Amaral MD, Dorfman R, Zielenski J, Masica DL, Karchin R, Millen L, Thomas PJ, Patrinos GP, Corey M, Lewis MH, Rommens JM, Castellani C, …, Cutting GR. Defining the disease liability of variants in the cystic fibrosis transmembrane conductance regulator gene. Nature Genetics 2013; 45: 1160–1167.

11. Clancy JP, Cotton CU, Donaldson SH, Solomon GM, VanDevanter DR, Boyle MP, Gentzsch M, Nick JA, Illek B, Wallenburg JC, Sorscher EJ, Amaral MD, Beekman JM, Naren AP, Bridges RJ, Thomas PJ, Cutting G, Rowe S, Durmowicz AG, …, Tuggle KL. CFTR modulator theratyping: Current status, gaps and future directions. J Cyst Fibros 2019; 18: 22–34.

12. Mannucci PM, Tuddenham EG. The hemophilias--from royal genes to gene therapy. N Engl J Med 2001; 344: 1773–1779.

13. Wettstein S, Underhaug J, Perez B, Marsden BD, Yue WW, Martinez A, Blau N. Linking genotypes database with locus-specific database and genotype-phenotype correlation in phenylketonuria. Eur J Hum Genet 2015; 23: 302–309.

14. Okano Y, Eisensmith RC, Güttler F, Lichter-Konecki U, Konecki DS, Trefz FK, Dasovich M, Wang T, Henriksen K, Lou H. Molecular basis of phenotypic heterogeneity in phenylketonuria. N Engl J Med 1991; 324: 1232–1238.

15. Jinnah HA, De Gregorio L, Harris JC, Nyhan WL, O’Neill JP. The spectrum of inherited mutations causing HPRT deficiency: 75 new cases and a review of 196 previously reported cases. Mutat Res 2000; 463: 309–326.

16. Taylor C, Commander CW, J.M. C, Strug LJ, Li W, Wright FA, Webel AD, Pace RG, Stonebraker JR, Naughton KM, Dorfman R, Sanford A, Blackman SM, Berthiaume Y, Pare P, Drumm ML, Zielenski J, Durie PR, Cutting GR, …, Corey M. A novel lung disease phenotype adjusted for mortality attrition for cystic fibrosis genetic modifier studies. Pediatric Pulmonology 2011; Epub no. doi: 10.1002/ppul.21456.

17. Han ST, Rab A, Pellicore MJ, Davis EF, McCague AF, Evans TA, Joynt AT, Lu Z, Cai Z, Raraigh KS, Hong JS, Sheppard DN, Sorscher EJ, Cutting GR. Residual function of cystic fibrosis mutants predicts response to small molecule CFTR modulators. JCI Insight 2018; 3.

18. Van Goor F, Yu H, Burton B, Hoffman BJ. Effect of ivacaftor on CFTR forms with missense mutations associated with defects in protein processing or function. J Cyst Fibros 2014; 13: 29–36.

19. Bihler H, Sivachenko A, Millen L, Bhatt P, Patel AT, Chin J, Bailey V, Musisi I, LaPan A, Allaire NE, Conte J, Simon NR, Magaret AS, Raraigh KS, Cutting GR, Skach WR, Bridges RJ, Thomas PJ, Mense M. In vitro modulator responsiveness of 655 CFTR variants found in people with cystic fibrosis. J Cyst Fibros 2024; 23: 664–675.

20. Welsh MJ, Smith AE. Molecular mechanisms of CFTR chloride channel dysfunction in cystic fibrosis. Cell 1993; 73: 1251–1254.

21. Burgel PR, Girodon E, Sharma N, Raynal C, Da Silva J, Sasorith S, Martin C, Sermet-Gaudelus I, Raraigh K. Elexacaftor-tezacaftor-ivacaftor in people with cystic fibrosis harbouring two. EClinicalMedicine 2025; 88: 103476.

22. Pion A, Kavanagh E, Joynt AT, Raraigh KS, Vanscoy L, Langfelder-Schwind E, McNamara J, Moore B, Patel S, Merlo K, Temme R, Capurro V, Pesce E, Merlo C, Pedemonte N, Cutting GR, Sharma N. Investigation of CFTR Function in Human Nasal Epithelial Cells Informs Personalized Medicine. Am J Respir Cell Mol Biol 2024; 71: 577–588.

23. Collaco JM, Blackman SM, Raraigh KS, Corvol H, Rommens JM, Pace RG, Boelle PY, McGready J, Sosnay PR, Strug LJ, Knowles MR, Cutting GR. Sources of Variation in Sweat Chloride Measurements in Cystic Fibrosis. Am J Respir Crit Care Med 2016; 194: 1375–1382.

24. Raraigh KS, Han ST, Davis E, Evans TA, Pellicore MJ, McCague AF, Joynt AT, Lu Z, Atalar M, Sharma N, Sheridan MB, Sosnay PR, Cutting GR. Functional Assays Are Essential for Interpretation of Missense Variants Associated with Variable Expressivity. Am J Hum Genet 2018; 102: 1062–1077.

25. Nykamp K, Truty R, Riethmaier D, Wilkinson J, Bristow SL, Aguilar S, Neitzel D, Faulkner N, Aradhya S. Elucidating clinical phenotypic variability associated with the polyT tract and TG repeats in CFTR. Hum Mutat 2021; 42: 1165–1172.

26. Wine JJ. How the sweat gland reveals levels of CFTR activity. J Cyst Fibros 2022; 21: 396–406.

27. Raju SV, Jackson PL, Courville CA, McNicholas CM, Sloane PA, Sabbatini G, Tidwell S, Tang LP, Liu B, Fortenberry JA, Jones CW, Boydston JA, Clancy JP, Bowen LE, Accurso FJ, Blalock JE, Dransfield MT, Rowe SM. Cigarette smoke induces systemic defects in cystic fibrosis transmembrane conductance regulator function. Am J Respir Crit Care Med 2013; 188: 1321–1330.

28. Burgel PR, Da Silva J, Girodon E, Durieu I, Reynaud-Gaubert M, Murris-Espin M, Chiron R, Grenet D, Ramel S, Mely L, Hamidfar R, Douvry B, Houdouin V, Audousset C, Macey J, Mittaine M, Weiss L, Cosson L, Danner-Boucher I, …, group FCFRNs. Sweat chloride and lung function responses to elexacaftor-tezacaftor-ivacaftor in people with cystic fibrosis with two versus one responsive CFTR variants: an analysis of two real-world observational studies. Lancet Respir Med 2025; 13: 978–989.

29. Vanscoy LL, Blackman SM, Collaco JM, Bowers A, Lai T, Naughton K, Algire M, McWilliams R, Beck S, Hoover-Fong J, Hamosh A, Cutler D, Cutting GR. Heritability of lung disease severity in cystic fibrosis. Am J Respir Crit Care Med 2007; 175: 1036–1043.

30. Mention K, Cavusoglu-Doran K, Joynt AT, Santos L, Sanz D, Eastman AC, Merlo C, Langfelder-Schwind E, Scallan MF, Farinha CM, Cutting GR, Sharma N, Harrison PT. Use of adenine base editing and homology-independent targeted integration strategies to correct the cystic fibrosis causing variant, W1282X. Hum Mol Genet 2023; 32: 3237–3248.

31. Vaidyanathan S, Baik R, Chen L, Bravo DT, Suarez CJ, Abazari SM, Salahudeen AA, Dudek AM, Teran CA, Davis TH, Lee CM, Bao G, Randell SH, Artandi SE, Wine JJ, Kuo CJ, Desai TJ, Nayak JV, Sellers ZM, Porteus MH. Targeted replacement of full-length CFTR in human airway stem cells by CRISPR-Cas9 for pan-mutation correction in the endogenous locus. Mol Ther 2022; 30: 223–237.

32. Kavanagh EW, Joynt AT, Pion AR, Eastman AC, Parr AI, Starego KL, Jain M, Shannon SR, Yoo EJ, Newby GA, Tzeng SY, Sharma N, Green JJ, Cutting GR. Base editing and nanoparticle transfection of airway cell types essential for treatment of cystic fibrosis. JCI Insight 2026; 11.

33. Mayer-Hamblett N, Clancy JP, Jain R, Donaldson SH, Fajac I, Goss CH, Polineni D, Ratjen F, Quon BS, Zemanick ET, Bell SC, Davies JC, Jain M, Konstan MW, Kerper NR, LaRosa T, Mall MA, McKone E, Pearson K, …, Downey DG. Advancing the pipeline of cystic fibrosis clinical trials: a new roadmap with a global trial network perspective. Lancet Respir Med 2023; 11: 932–944.

