## Supplementary figures and images for "Clinical and primary cell evidence reveals complex CFTR function–phenotype relationships"

### Supplemental Figure 1

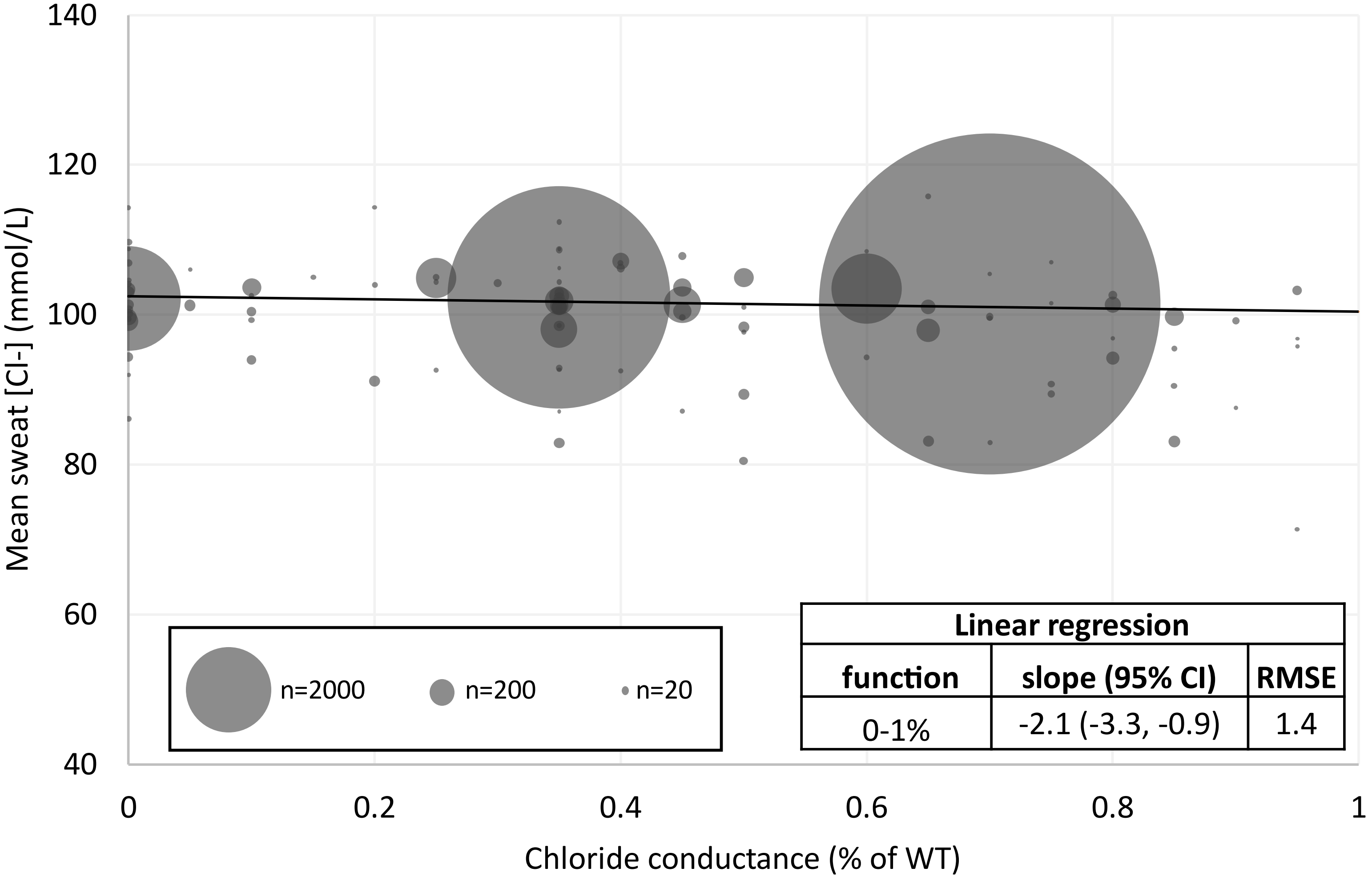
